# Immune–metabolic PET/MRI uncovers microenvironmental reprogramming under combined immunotherapy and anti-angiogenic therapy

**DOI:** 10.64898/2026.08.06.743278

**Authors:** Sixing Li, Marie-Aline Neveu, Laura Kuebler, Stefania Pezzana, Ainara Barco-Tejada, Ian Wilson, Irene Gonzalez-Menendez, Leticia Quintanilla-Martínez, Dominik Sonanini, Andreas M. Schmid, Manfred Kneilling, André F. Martins

## Abstract

The limited efficacy of immune checkpoint inhibitor (ICI) therapy in triple-negative breast cancer (TNBC) highlights the need for combination strategies that enhance antitumor responses. Sorafenib, a multikinase inhibitor with anti-angiogenic and immunomodulatory activity, represents a rational partner for ICI-based combination therapy. However, therapeutic responses to such combinations are biologically complex and cannot be fully characterized by any single biomarker or imaging modality. Here, we evaluated ICI therapy combined with sorafenib in the aggressive and ICI-refractory orthotopic 4T1 TNBC model. Therapeutic responses were assessed using a unique longitudinal multimodal imaging framework integrating [⁸⁹Zr]Zr-DFO–anti-CD8 minibody and [¹⁸F]FDG PET, as well as perfluorocarbon (PFC)-based ¹⁹F MRI and hyperpolarized ¹³C MRS, together with *ex vivo* analyses. Only the ICI–sorafenib combination suppressed tumor growth, whereas both monotherapies showed limited antitumor activity. Multimodal imaging, together with complementary *ex vivo* analyses, uncovered coordinated tumor microenvironment (TME) remodeling, including vascular normalization, elevated CD8⁺ cell presence with modest enrichment in the tumor center, delayed increase in phagocyte-associated ¹⁹F MRI signal coupled with reduced CD206⁺ cell infiltration, and sustained metabolic activity. These findings support ICI–sorafenib combination therapy as a promising therapeutic strategy for TNBC. Therapeutic efficacy reflected coordinated vascular, immune, and metabolic remodeling. This multimodal imaging framework enables non-invasive longitudinal monitoring of these complementary TME changes, providing a comprehensive strategy for treatment assessment in immunotherapy-based combination therapies.

**One Sentence Summary:** Longitudinal multimodal imaging identified a multidimensional TME response signature of effective ICI-sorafenib therapy in TNBC.

## INTRODUCTION

Female breast cancer was the second most commonly diagnosed cancer in 2022 and the fourth leading cause of cancer-related death worldwide (*1*). Triple-negative breast cancer (TNBC) accounts for 15% to 20% of all breast cancer and is associated with particularly poor clinical outcomes (*2, 3*). Among all breast cancer subtypes, TNBC is considered the most immunogenic (*4*) and has emerged as a major focus of immune checkpoint inhibitor (ICI) therapy. However, only a limited proportion of TNBC patients respond to ICI treatment, despite durable clinical benefit in a subset of patients (*5, 6*). Chemotherapy continues to dominate first-line treatment (*7*), while emerging approaches such as antibody–drug conjugates continue to expand therapeutic options (*8*). In parallel, combination strategies aimed at enhancing responses to ICIs remain an active area of investigation.

Among potential ICI combination compounds, anti-angiogenic therapy has attracted particular interest in TNBC (*9*). Preclinical studies using VEGFR2 blockade or nintedanib, a tyrosine kinase inhibitor (TKI) targeting VEGFR, FGFR, and PDGFR, have shown that anti-angiogenic treatment can enhance the efficacy of ICI therapy and remodel the tumor microenvironment (TME), potentially facilitating immune-cell entry into tumors (*10, 11*). Results from the single-arm ATRACTIB phase II trial, which evaluated an anti-PD-L1 monoclonal antibody (mAb) combined with an anti-VEGF mAb and paclitaxel, further support the clinical rationale for combining ICI therapy with anti-angiogenic therapy in advanced TNBC (*9*).

Despite encouraging results, most studies have focused on VEGF/VEGFR-targeted therapies and a limited subset of anti-angiogenic TKIs, leaving multikinase inhibitors largely unexplored in this setting. Sorafenib, an FDA-approved multikinase inhibitor targeting VEGFR, PDGFR, and RAF (*12, 13*), has shown promising activity in combination with ICI in hepatocellular carcinoma (HCC) in preclinical cancer studies (*14*) and a prospective phase II clinical trial (*15*). However, this therapeutic strategy has not been systematically investigated in TNBC.

Although tumor growth remains the primary measure of therapeutic efficacy (*16*), it provides limited insight into the dynamic biological processes that precede and shape treatment response within the TME. Multimodal imaging provides a framework for longitudinal monitoring of treatment-induced alterations within the TME. CD8 PET enables assessment of treatment-induced changes in CD8⁺ T-cell infiltration and has been successfully applied in breast cancer models receiving ICOS agonist alone or in combination with anti-PD-1 therapy (*17*). Complementarily, perfluorocarbon (PFC)-based ¹⁹F MRI enables non-invasive tracking of phagocyte infiltration, including tumor-associated macrophages (TAMs), and has been used to monitor therapy-induced myeloid cell dynamics in breast cancer and other solid tumors (*18, 19*). Metabolic imaging further extends this assessment by providing functional information. [¹⁸F]FDG PET has been widely used to assess treatment-induced metabolic changes during ICI therapy (*20, 21*). However, its interpretation may be confounded by pseudoprogression, as immune activation markedly increases glucose metabolism in activated T cells and myeloid cells in addition to tumor cells (*22, 23*). Hyperpolarized [1-¹³C]pyruvate MRS (HP MRS), which measures pyruvate-to-lactate conversion, provides insight into downstream glycolytic metabolism and has detected therapy-induced metabolic changes in preclinical melanoma models (*24*), but has seen only limited application in breast cancer. Despite the complementary strengths of immune and metabolic imaging, their integration for longitudinal assessment of ICI-based combination therapies has not been systematically investigated.

In this study, we used the orthotopic 4T1 TNBC model to evaluate dual anti-PD-1/CTLA-4 blockade combined with sorafenib. The 4T1 model exhibits substantial immune-cell infiltration but remains largely refractory to ICI or sorafenib monotherapy (*25, 26*), providing a stringent setting to evaluate combination therapy. We assessed therapeutic response by longitudinal multimodal imaging using [⁸⁹Zr]Zr-DFO–anti-CD8 minibody (Mb) PET, PFC-based ¹⁹F MRI, [¹⁸F]FDG PET, and hyperpolarized [1-¹³C]pyruvate MRS (HP MRS) to monitor CD8⁺ T-cell infiltration, phagocyte-associated changes, glucose uptake, and pyruvate-to-lactate conversion, respectively. Imaging findings were validated by complementary *ex vivo* analyses. This integrated approach enables longitudinal, multidimensional characterization of therapeutic response beyond any single biomarker or imaging modality.

## RESULTS

### Combined ICI and sorafenib treatment improves antitumor efficacy in mice with orthotopic 4T1 tumors

To evaluate therapeutic efficacy, 4T1 tumor-bearing mice received dual anti-PD-1/CTLA-4 mAbs plus sorafenib (Combo), dual anti-PD-1/CTLA-4 mAbs alone (ICIs), sorafenib alone (Sorafenib), or isotype control (Control). To capture the spatial temporal dynamics of the treatment response, multimodal imaging was performed longitudinally at baseline (TP0), at an early time point during therapy (TP1, D8), and at a late time point during therapy (TP2, D12-15) (Fig. 1A). [⁸⁹Zr]Zr-DFO– anti-CD8 Mb PET was performed separately using a single-tracer administration followed by serial imaging throughout the first week of therapy. Tumor growth analysis showed efficient suppression of 4T1 tumor progression only in the Combo treatment group (Combo versus Control, *p*<0.0001), whereas neither ICIs nor Sorafenib monotherapy exhibited a measurable therapeutic benefit (Fig. 1B). Consistently, endpoint (TP2) tumor weight analysis confirmed a significant reduction in tumor burden exclusively in the Combo treatment group (Fig. 1C). Importantly, body weight remained stable in all experimental groups throughout the study, supporting the favorable tolerability of the treatment regimens (Fig. 1D). These results demonstrate that only Combo treatment achieved antitumor efficacy in the 4T1 TNBC model.

**Fig. 1.**
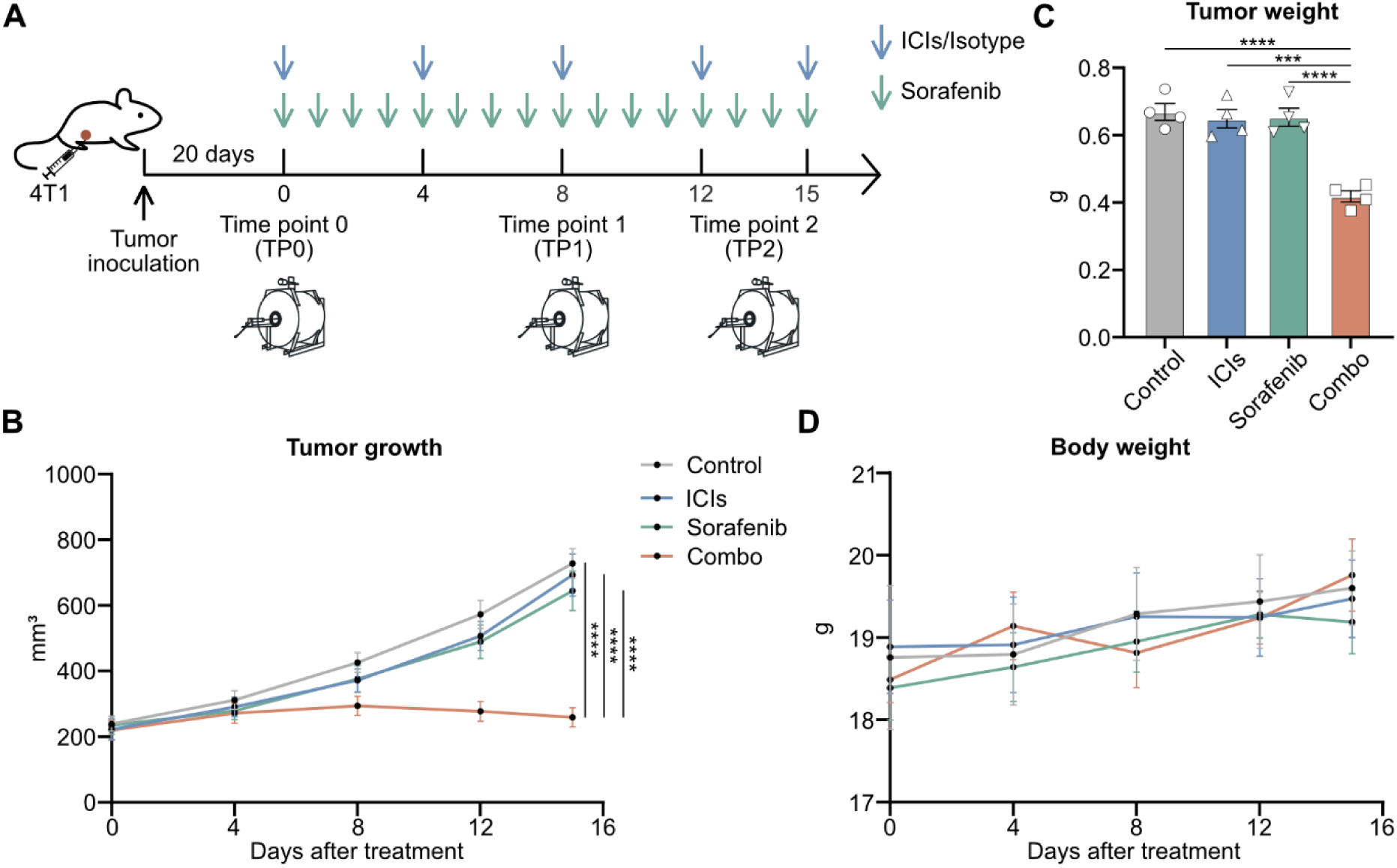
Experimental timeline and antitumor effect of immune checkpoint inhibitors (ICIs) combined with sorafenib *in vivo*. **(A)** Schematic illustration of the experimental timeline, including tumor cell inoculation, treatment administration, imaging time points (TP0, TP1, and TP2). **(B)** Longitudinal tumor growth during the treatment period. **(C)** Tumor weights at TP2. **(D)** Changes in body weight throughout the treatment period. For panels B to D, n=3–5 mice per group. Data are shown as mean±SEM. Each dot represents one mouse. Statistical analyses were performed using repeated-measures two-way ANOVA with Tukey’s test or Brown-Forsythe and Welch ANOVA with Dunnett’s T3 test, as appropriate. \*\*\**p*<0.001, \*\*\*\**p*<0.0001.

### Combo treatment is associated with increased tumor vascular maturity

Given the anti-angiogenic activity of sorafenib, we next examined treatment-induced vascular changes that may influence drug delivery and immune infiltration. Tumors collected at TP2 were analyzed by CD31 and α-SMA immunohistochemistry (IHC), revealing distinct vascular patterns across treatment groups (Fig. 2A and fig. S1). Tumors from Control-treated mice displayed a high density of CD31⁺ vessels with limited α-SMA⁺ pericyte coverage, consistent with an immature vascular phenotype. Tumors from ICIs-treated mice showed similar vessel density but increased α-SMA⁺ pericyte coverage. In contrast, tumors from Sorafenib- and Combo-treated mice exhibited markedly reduced CD31⁺ vessel density with preserved or prominent α-SMA⁺ pericyte coverage. Quantification confirmed significantly lower CD31⁺ vessel density in the Sorafenib and Combo groups compared to the Control and ICIs groups (Fig. 2B). Vascular maturation in the TME, assessed by the spatial overlap of CD31⁺ vessels and α-SMA⁺ pericytes, was highest in the Combo-treated group, intermediate in the ICIs and Sorafenib treatment groups, and lowest in Control group (Fig. 2C). Together, these findings indicate that sorafenib reduced vessel density, ICIs enhanced pericyte coverage, and their combination yielded the most mature vascular phenotype in the TME.

**Fig. 2.**
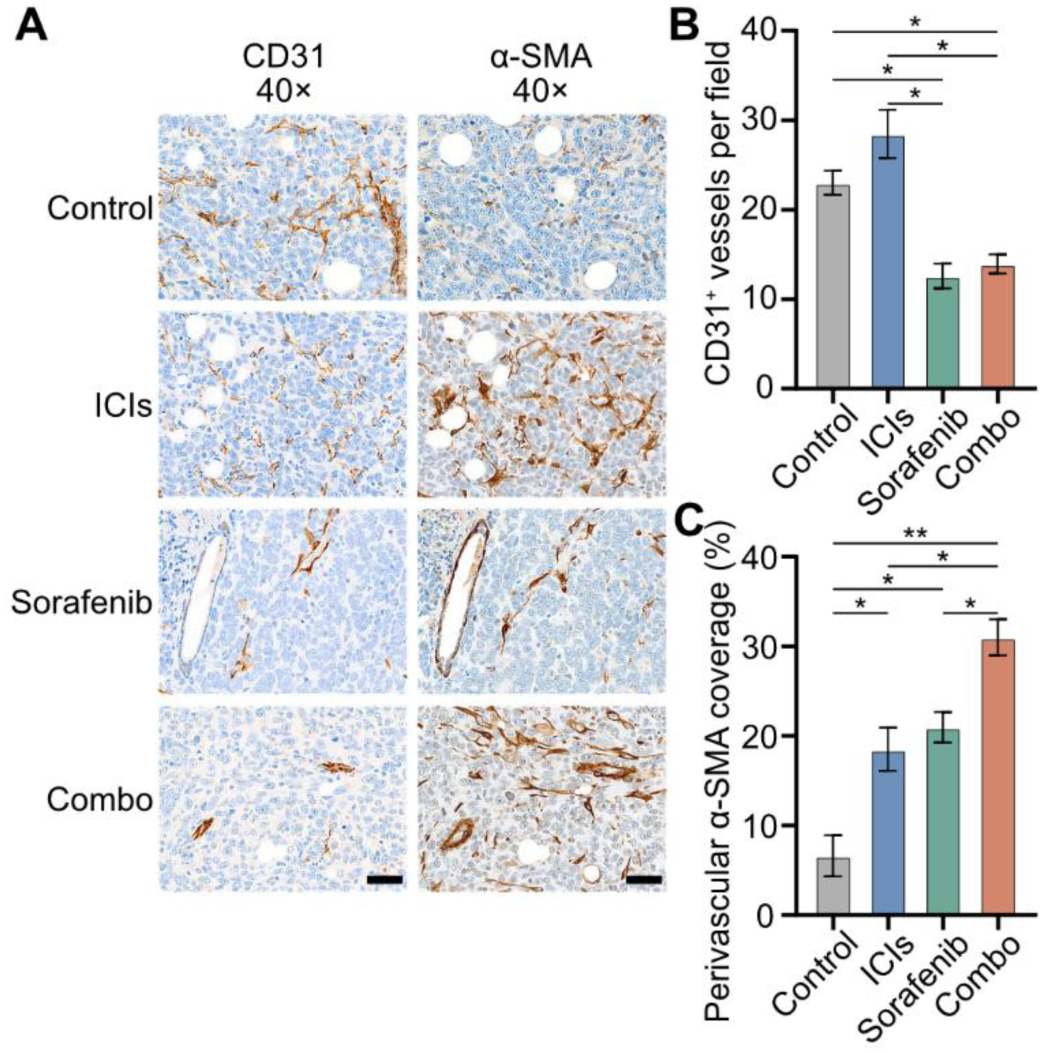
Treatment-induced vascular changes. **(A)** Representative CD31 and α-SMA IHC images from serial tumor sections in matched 40× fields. Scale bars: 50µm. **(B)** CD31⁺ vessel counts per high-power field. **(C)** α-SMA⁺CD31⁺ area as percentage of CD31⁺ vessel area. For panels B and C, n=3–4 mice per group. Data are shown as mean±SEM. Statistical analyses were performed using Brown-Forsythe and Welch ANOVA with Dunnett’s T3 test. \**p*<0.05, \*\**p*<0.01.

### Combo treatment increases intratumoral CD8⁺ cell infiltration and distribution

Given the established link between tumor vasculature and immune-cell trafficking, we next examined whether treatment altered intratumoral CD8⁺ cell infiltration. To longitudinally assess CD8⁺ cell dynamics, mice received a single injection of [⁸⁹Zr]Zr-DFO–anti-CD8 Mb one day before treatment initiation and underwent serial PET/MRI at baseline (D0) and on D1, D3, and D7 after treatment initiation (Fig. 3A). Representative [⁸⁹Zr]Zr-DFO–anti-CD8 Mb PET images co-registered with T2-weighted MRI showed a reduced intratumoral tracer uptake over time, with treatment-related differences emerging at later imaging time points (Fig. 3B). At D3, quantification of mean intratumoral [⁸⁹Zr]Zr-DFO–anti-CD8 Mb uptake confirmed a significantly enhanced uptake in tumors of the Combo treatment group when compared to the tumors of the Control (*p*=0.001) and ICIs (*p*=0.01) treatment groups. At D7, the tracer uptake in the tumors of the Combo treatment group was significantly enhanced when compared to tumors of all other treatment groups (Fig. 3C, *p*<0.05).

**Fig. 3.**
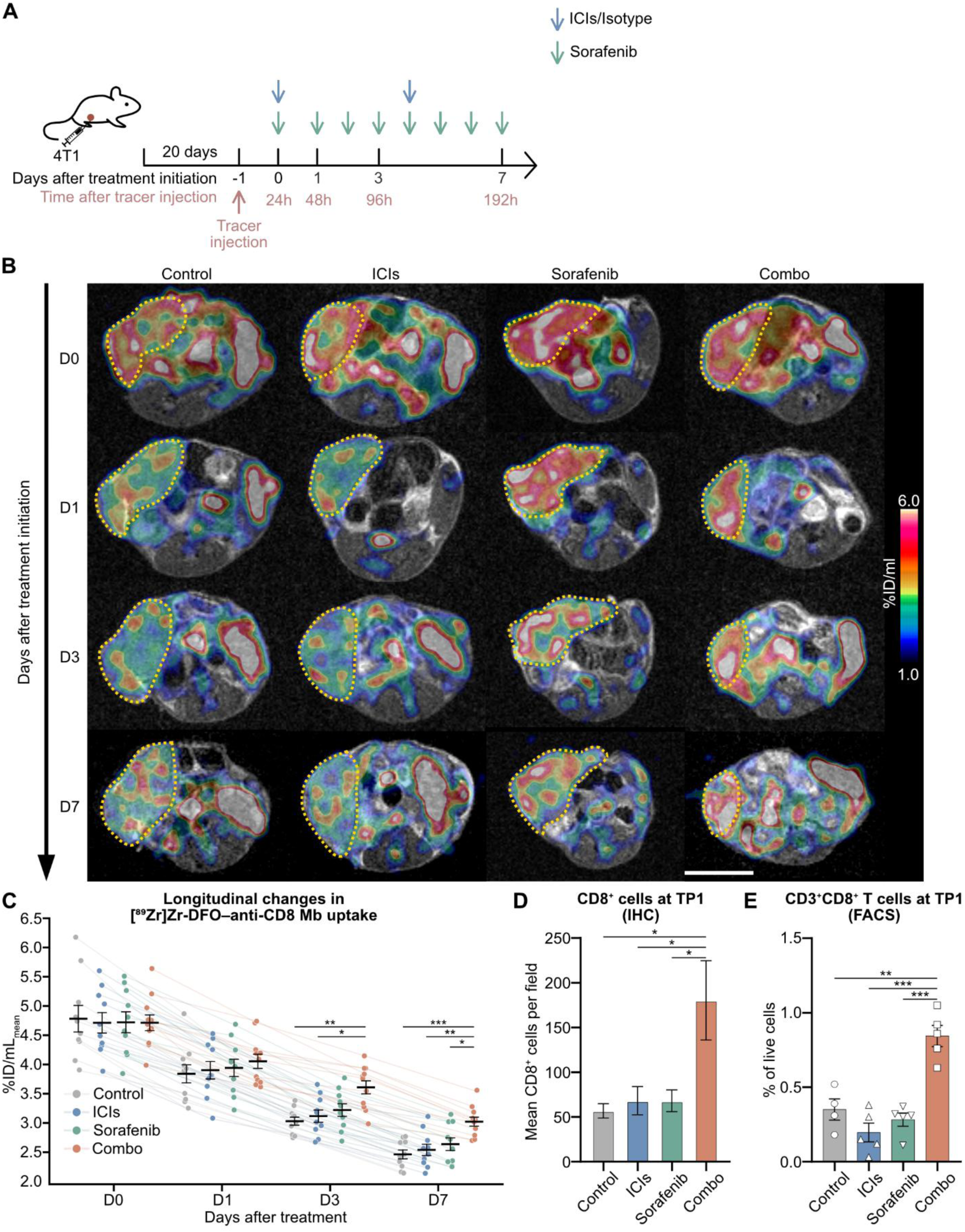
Tracking CD8-targeted PET signal and T-cell infiltration during treatment. **(A)** Schematic illustration of the [⁸⁹Zr]Zr-DFO–anti-CD8 minibody (Mb) PET imaging protocol. **(B)** Representative [⁸⁹Zr]Zr-DFO–anti-CD8 Mb PET/MRI axial slices from individual animals in each treatment group across D0, D1, D3, and D7. Scale bar: 1cm. **(C)** Longitudinal quantification of [⁸⁹Zr]Zr-DFO–anti-CD8 Mb uptake as %ID/mL. **(D)** Quantification of CD8⁺ cell counts by IHC. **(E)** Quantification of CD3^+^CD8⁺ T cells as percentage of live cells by flow cytometry (FACS). For panel C, n=10–11 mice per group; for panels D and E, n=3–5 mice per group. Data are shown as mean±SEM. Each dot represents one mouse. Statistical analyses were performed using repeated-measures two-way ANOVA with Tukey’s test or Brown-Forsythe and Welch ANOVA with Dunnett’s T3 test, as appropriate. \**p*<0.05, \*\**p*<0.01, \*\*\**p*<0.001.

To support the imaging findings, mice were sacrificed one day after the final [⁸⁹Zr]Zr-DFO–anti-CD8 Mb PET/MRI scan and tumors were collected for *ex vivo* analysis. Quantification of CD8⁺ cells by IHC demonstrated significantly higher intratumoral CD8⁺ cell infiltration in the Combo group than in the other treatment groups, reaching up to approximately threefold higher levels (Fig. 3D, p < 0.05). These results were further supported by flow cytometry analysis of tumors, revealing a significantly enhanced frequency of CD3^+^CD8⁺ T cells in tumors of the Combo treatment group, when compared to the other treatment groups (Fig. 3E, *p*<0.01). In summary, our *ex vivo* analysis suggests the increase in [⁸⁹Zr]Zr-DFO–anti-CD8 Mb uptake in the Combo treatment group is associated to enhanced intratumoral expression of CD8⁺ T cells.

We next investigated whether Combo treatment altered the intratumoral distribution of CD8⁺ cells. Longitudinal [⁸⁹Zr]Zr-DFO–anti-CD8 Mb PET images were used to assess their spatial distribution by generating radial uptake profiles (Fig. 4, A and B). At D0, all groups exhibited similar profiles, with higher tracer uptake at the tumor periphery. By D7, however, intratumoral tracer uptake in the Combo treatment group shifted towards the tumor center. Accordingly, the center-to-periphery [⁸⁹Zr]Zr-DFO–anti-CD8 Mb uptake ratio was comparable across groups at baseline but increased exclusively in tumors of the Combo treatment group at D7 (Fig. 4C, *p*<0.01). To validate these findings, the corresponding CD8 IHC sections of tumors were analyzed for the spatial distribution of CD8⁺ cells (Fig. 4D and fig. S3). Despite differences in the definitions of central and peripheral regions between PET and IHC analyses, tumors from Combo-treated mice displayed a significantly higher center-to-periphery CD8⁺ cell ratio than those from all other treatment groups (Fig. 4E, *p*<0.05). Together, these findings demonstrate that Combo treatment promotes infiltration of CD8⁺ cells towards the tumor core.

**Fig. 4.**
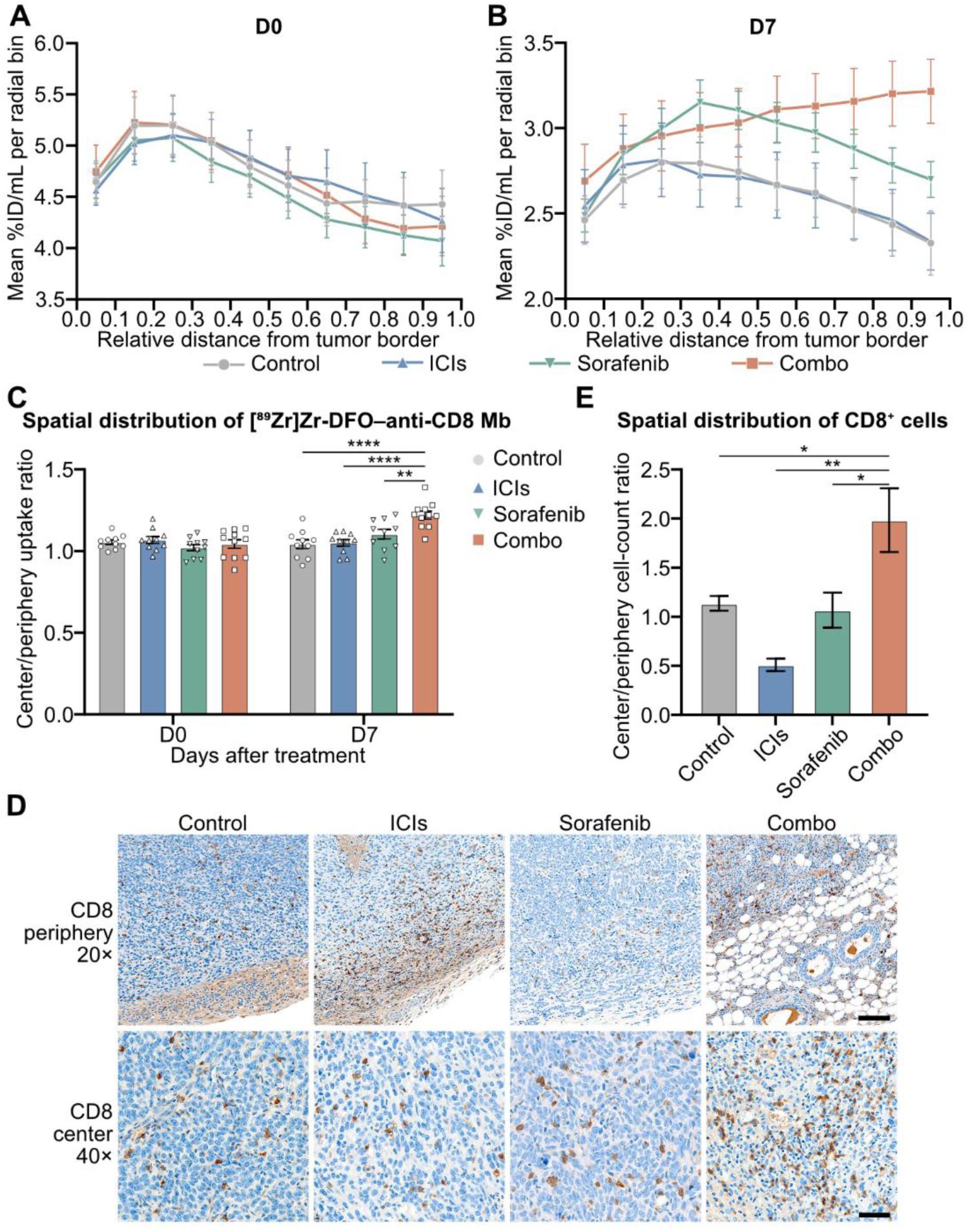
Spatial redistribution of [⁸⁹Zr]Zr-DFO–anti-CD8 Mb and CD8⁺ cells during treatment. (A–B) Radial profiles of [⁸⁹Zr]Zr-DFO–anti-CD8 Mb uptake at D0 (A) and D7 (B). The x-axis represents the normalized radial distance from the tumor border (0) to the tumor core (1). **(C)** Center/periphery ratio of [⁸⁹Zr]Zr-DFO–anti-CD8 Mb uptake at D0 and D7. **(D)** Representative CD8 IHC images showing tumor periphery and center regions at TP1. Scale bars: 100µm for periphery images (20×) and 50µm for center images (40×). **(E)** Quantification of center/periphery CD8⁺ cell count ratio by IHC at TP1. For panels A to B, n=10–11 mice per group; for panel E, n=3 mice per group. Data are shown as mean±SEM. Each dot represents one mouse. Statistical analyses were performed using repeated-measures two-way ANOVA with Tukey’s test or one-way ANOVA with Dunnett’s test, as appropriate. \**p*<0.05, \*\**p*<0.01, \*\*\*\**p*<0.0001.

### Combo treatment enhances intratumoral phagocyte recruitment and reduced infiltration of CD206**^+^**TAMs

We next examined treatment-induced phagocyte recruitment by PFC-based ¹⁹F MRI. Representative ¹⁹F/¹H MRI revealed low intratumoral ¹⁹F signal at TP0 and TP1 across all treatment groups, whereas the ¹⁹F signal increased selectively in the Combo treatment group at TP2 (Fig. 5A), indicating enhanced phagocyte accumulation. Quantitative analysis of the ^19^F signal confirmed these observations. The intratumoral ¹⁹F signal remained low until TP1 but increased significantly at TP2 exclusively in the Combo treatment group (Fig. 5B, *p*<0.05).

**Fig. 5.**
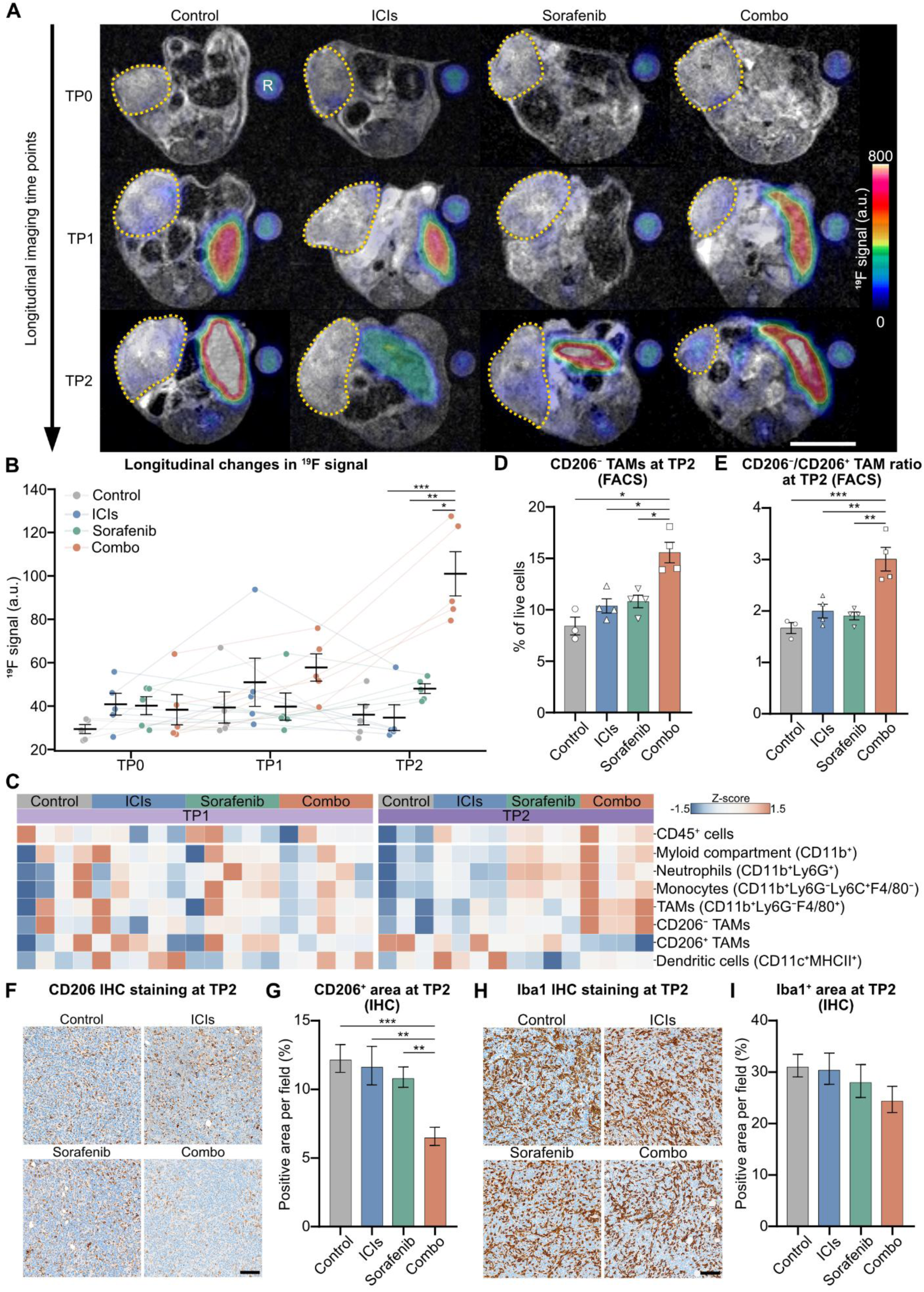
¹⁹F MRI signal and CD206-associated phenotype shift following treatment. **(A)** Representative ¹⁹F/¹H MRI axial slices from individual animals in each treatment group across TP0, TP1, and TP2. Scale bar: 1cm. R indicates the ^19^F reference tube. **(B)** Longitudinal quantification of ¹⁹F MRI signal intensity. **(C)** Heatmaps of flow cytometry data showing myeloid and phagocyte-associated populations at TP1 and TP2. **(D)** Quantification of CD206⁻ TAMs as percentage of live cells at TP2 by flow cytometry. **(E)** Quantification of CD206⁻/CD206⁺ TAM ratio at TP2 by flow cytometry. **(F–I)** Representative CD206 (F) and Iba1 (H) IHC images with quantification of CD206⁺ (G) and Iba1⁺ (I) area at TP2. Scale bars: 50µm. Cell populations were defined according to the gating strategy presented in fig. S2, A and B. For quantitative panels, n=3–5 mice per group. Each dot or column represents one mouse. Quantitative data are shown as mean±SEM. Statistical analyses were performed using repeated-measures two-way ANOVA with Tukey’s test, Brown-Forsythe and Welch ANOVA with Dunnett’s T3 test, or one-way ANOVA with Dunnett’s test, as appropriate. \**p*<0.05, \*\**p*<0.01, \*\*\**p*<0.001.

For *ex vivo* cross correlation of the PFC-based ¹⁹F MRI data, we performed flow cytometry (TP1 and TP2) and IHC (TP2) of tumors collected from mice of all treatment groups and focused on myeloid cell expression and differentiation patterns. Flow cytometry analysis of all treatment groups (n=4-5) focused on the myeloid compartment (CD45^+^CD11b^+^) in the tumors, especially on neutrophils (CD45^+^CD11b^+^Ly6G^+^), monocytes (CD45^+^CD11b^+^Ly6G⁻Ly6C^+^F4/80^⁻^), TAMs (CD45^+^CD11b^+^Ly6G⁻F4/80^+^), CD206⁻ TAMs (CD45^+^CD11b^+^Ly6G⁻F4/80^+^CD206⁻), CD206^+^ TAMs (CD45^+^CD11b^+^Ly6G⁻F4/80^+^CD206^+^), as well as dendritic cells (CD45^+^CD11c^+^MHCII^+^). Generally, we observed a high degree of variation within the treatment groups (Fig. 5C). At TP1, we observed no significant differences among all treatment groups, consistent with the low ¹⁹F MRI signal at this time point. At TP2, however, the Combo treatment group showed the most prominent increase in the myeloid compartment (Fig. 5C). We determined a significant increase in CD206⁻ TAMs (Fig. 5D, *p*<0.05) and in the CD206⁻/CD206^+^ TAM ratio (Fig. 5E, *p*<0.01) when compared to all the other treatment groups.

To further characterize TAM phenotypes, we additionally performed Iba1 and CD206 IHC of tumors collected at TP2. In alignment with our flow cytometry data, we observed a significant decrease in CD206⁺ TAMs in tumors of the Combo treatment group when compared to the tumors of all other treatment groups (Fig. 5, F and G, *p*<0.01). Focusing on phagocytes in the TME, we observed moderate but not significant decrease in the Combo treatment group when compared to the other treatment cohorts (Fig. 5, H and I).

Together, these findings suggest that Combo treatment remodels the myeloid compartment in 4T1 tumors, as reflected by a delayed increase in the phagocyte-associated ^19^F MRI signal and reduced infiltration of CD206⁺ TAMs at TP2.

### Metabolic imaging of tumors reveals a unique response to Combo treatment

We next examined treatment-associated changes in glucose metabolism in the TME of 4T1 tumors using [¹⁸F]FDG PET and HP MRS. Representative [¹⁸F]FDG PET/MRI images revealed heterogeneous intratumoral [¹⁸F]FDG uptake across all treatment groups and imaging time points (Fig. 6A). We observed that the mean tumor [¹⁸F]FDG uptake remained relatively stable in the ICIs and Combo treatment groups at all imaging time points. However, at TP2, we identified a significant decrease in the mean tumor [¹⁸F]FDG uptake in the Sorafenib and Control treatment groups (Fig. 6B). To further evaluate whether whole-tumor quantification influenced the interpretation of [¹⁸F]FDG PET findings, additional analyses using maximum [¹⁸F]FDG uptake and metabolic tumor volume (MTV) were performed. Maximum [¹⁸F]FDG uptake showed a broadly similar temporal pattern to mean uptake (fig. S4A). MTV analyses using 30% and 40% thresholds also showed broadly similar temporal pattern with a main exception at TP1 for the Combo and Control groups (fig. S4, B to C).

**Fig. 6.**
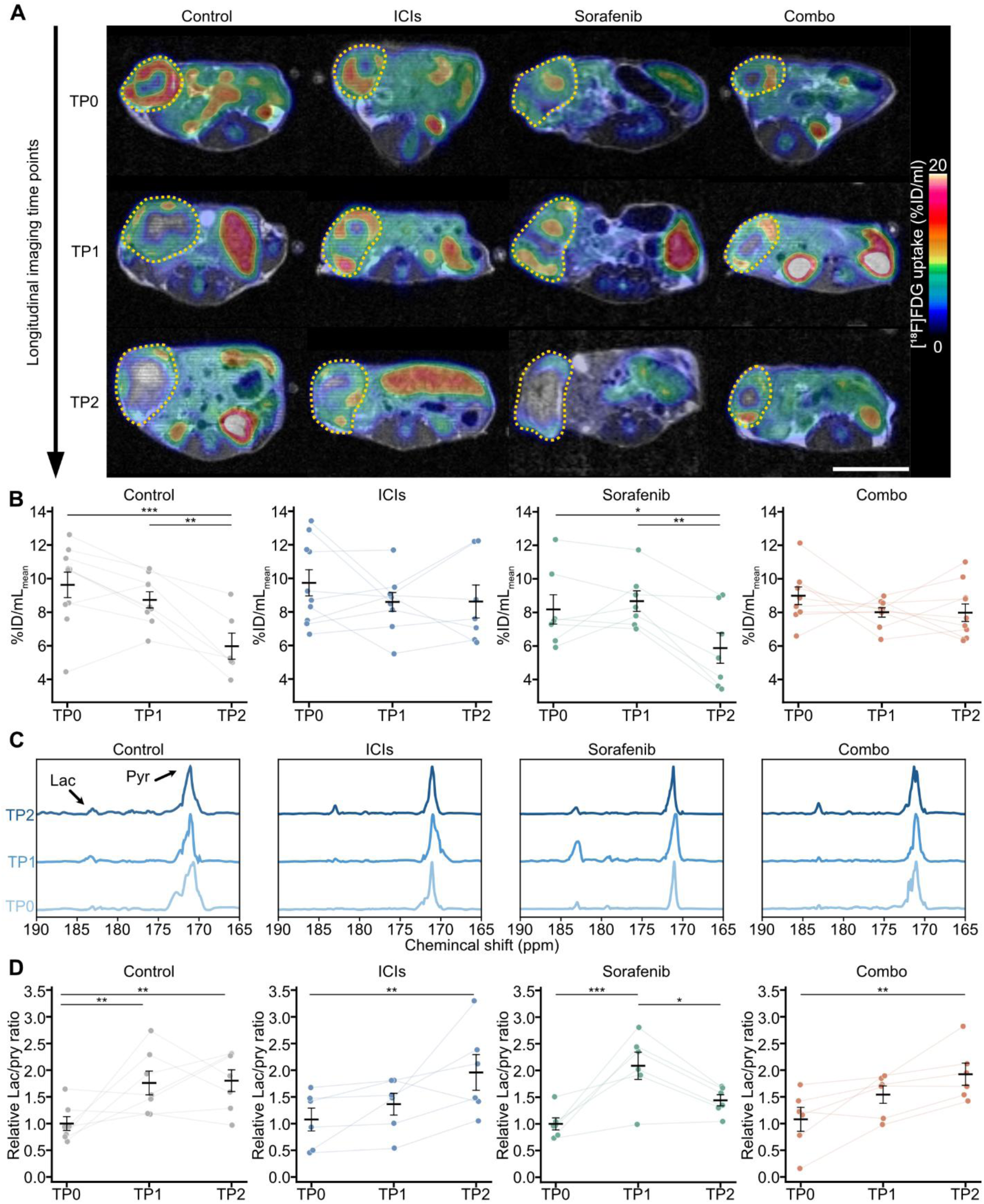
[¹⁸F]FDG PET and hyperpolarized [1-¹³C]pyruvate MRS (HP MRS) reveal treatment-associated metabolic changes. **(A)** Representative [¹⁸F]FDG PET/MRI axial slices from individual animals in each treatment group across TP0, TP1, and TP2. Scale bar: 1cm. **(B)** Longitudinal quantification of [¹⁸F]FDG uptake as %ID/mL within each treatment group. **(C)** Representative 2D HP MRS spectra across TP0, TP1, and TP2, showing pyruvate (Pyr) and lactate (Lac) peaks. (D) Longitudinal quantification of normalized Lac/Pyr ratio within each treatment group. Lac/Pyr ratios were normalized to the mean TP0 value of each treatment group. For quantitative panels, n=6–10 mice per group. Each dot represents one mouse. Data are shown as mean±SEM. Statistical analyses were performed using mixed-effects analysis followed by Tukey’s multiple-comparisons test. \**p*<0.05, \*\**p*<0.01, \*\*\**p*<0.001.

Longitudinal HP MRS spectra analysis of tumors displayed dynamic variations of lactate (Lac) and pyruvate (Pyr) signals across all treatment groups and imaging time points (Fig. 6C and fig. S3D). We observed a gradual increase of the relative Lac/Pyr ratio from TP0 to TP2 in tumors of Combo- or ICIs-treated mice. Tumors of the Sorafenib treatment group showed a transient increase in the relative Lac/Pyr ratio at TP1 followed by a decrease to baseline levels at TP2 (Fig. 6D). In tumors of the Control treatment group, the relative Lac/Pyr ratio increased from TP0 to TP1 and decreased slightly at TP2.

Together, these data identify a unique metabolic response to Combo treatment, with stable tumor glucose uptake despite progressively increasing pyruvate-to-lactate conversion during therapy.

### Integrated multimodal profiling reveals coordinated TME remodeling following Combo treatment

The multimodal design of this study enabled an integrated comparison of treatment-induced TME states across vascular, immune, and metabolic compartments. The radar plot summarizes seven representative parameters (Fig. 7). Overall, Combo treatment was associated with coordinated vascular, immune, and metabolic remodeling, highlighting the power of immune-metabolic multimodal imaging to resolve complex treatment response dynamics.

**Fig. 7.**
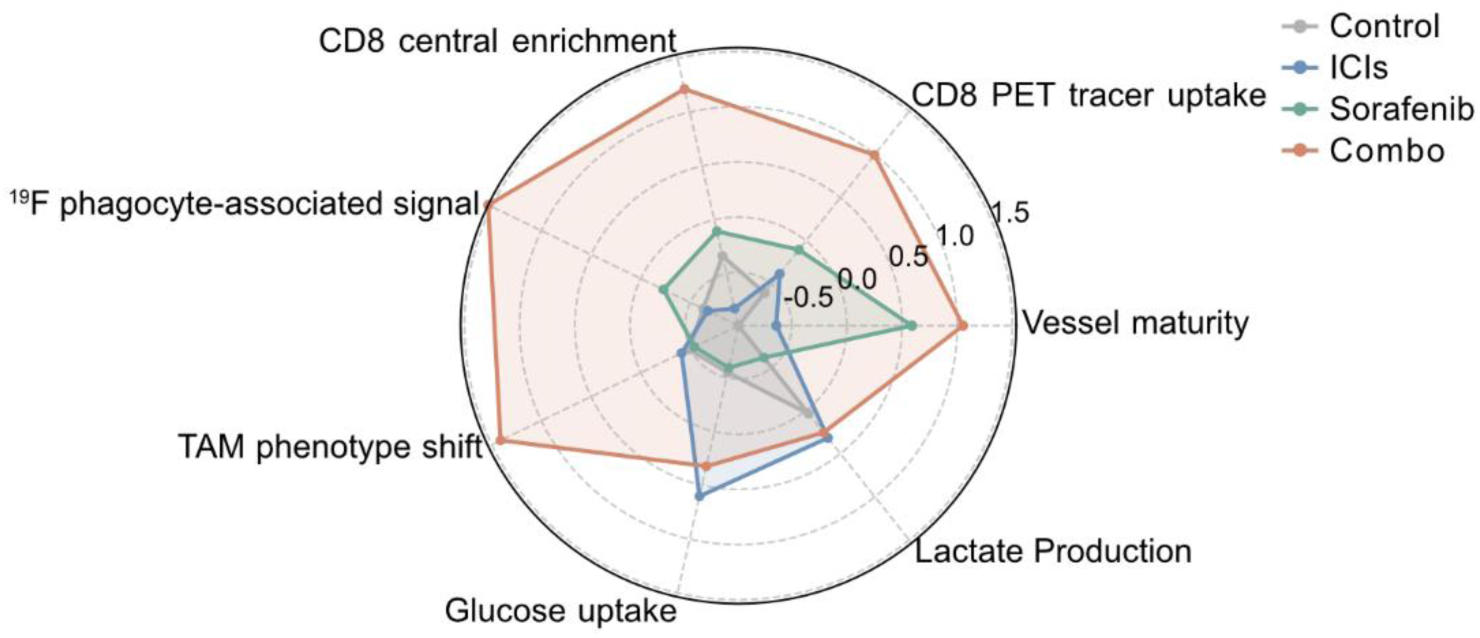
Summary of treatment-associated changes in the TME. The radar plot summarizes z-score-normalized vascular, immune, and metabolic parameters.

## DISCUSSION

Combination therapies with immune checkpoint inhibitors (ICIs), antibody-drug conjugates, and chemotherapies have entered first-line treatment in TNBC. In our study, the combination of ICI and sorafenib improved antitumor efficacy in the 4T1 model, whereas ICI and sorafenib monotherapies had limited effects. Longitudinal multimodal imaging and *ex vivo* analyses demonstrated coordinated remodeling of the TME. Combo treatment promoted vascular maturation, enhanced CD8⁺ cell infiltration with preferential accumulation in the tumor center, increased the phagocyte-associated ^19^F MRI signal, and reduced CD206⁺ TAM infiltration. Despite these changes, tumor glucose uptake remained stable while pyruvate-to-lactate conversion progressively increased.

Abnormal tumor vasculature limits immune-cell trafficking and contributes to an immunosuppressive TME (*27, 28*). Anti-angiogenic therapy has therefore been proposed to transiently normalize tumor vessels and improve immune infiltration, providing a rationale for combining anti-angiogenic agents with ICIs (*29, 30*). In our study, both monotherapies induced distinct aspects of vascular remodeling, consistent with previous reports (*31, 32*), but neither improved tumor control. In contrast, the Combo treatment combined these complementary vascular changes and achieved antitumor efficacy. A clinical study in patients with HCC showed that sorafenib alone reduced CD31⁺ microvessel density with limited tumor control, whereas sequential anti-PD-1 mAb treatment followed by sorafenib treatment preserved microvessel density, increased pericyte coverage, and achieved superior antitumor efficacy (*33*). Despite differences in tumor type and treatment schedule, our findings and those of the clinical study suggest that vascular remodeling alone does not necessarily predict therapeutic benefit. The reported CD8⁺ T cell dependence of these vascular and antitumor effects further highlights the close interplay between vascular remodeling and immune-cell infiltration during ICI–sorafenib combination therapy.

[⁸⁹Zr]Zr-DFO–anti-CD8 Mb PET enabled longitudinal assessment of both temporal and spatial changes in intratumoral CD8-associated tracer uptake during treatment. In our study, treatment-associated differences were detectable by [⁸⁹Zr]Zr-DFO–anti-CD8 Mb PET as early as D3, preceding major divergence in tumor growth. This early detection window is consistent with previous CD8 PET studies in breast cancer models, where treatment-induced increase in CD8 tracer uptake was detectable within 6–11 days after the onset of immunotherapy (*17, 34*). Notably, the early increase in [⁸⁹Zr]Zr-DFO–anti-CD8 Mb uptake observed in both sorafenib-treated groups may partly reflect sorafenib-mediated enhancement of tracer delivery, whereas only the Combo treatment group exhibited a further increase at D7. Previous MRI-based studies in the 4T1 model showed that sorafenib can induce a transient vascular normalization window, characterized by increased perfusion and permeability (*35*). PET studies with other tracers have also reported sorafenib-induced changes in tracer uptake, independent of the intended biological target, through altered vascular kinetics (*36*). Thus, the transient increase in intratumoral [⁸⁹Zr]Zr-DFO–anti-CD8 Mb uptake after sorafenib monotherapy may partly reflect improved tracer delivery due to vascular changes, whereas the sustained increase in the Combo treatment group is consistent with an enhanced CD8⁺ cell infiltration, as confirmed by flow cytometry and IHC. Unlike previous immunoPET distribution analysis that relied on qualitative pattern descriptions, line profiles, or regional peak counts (*17, 34, 37–39*), our radial profiling and center/periphery analysis provided a more objective and spatially defined characterization of [⁸⁹Zr]Zr-DFO–anti-CD8 Mb distribution patterns. By integrating early uptake changes with intratumoral spatial distribution, [⁸⁹Zr]Zr-DFO– anti-CD8 Mb PET provided a comprehensive longitudinal assessment of the adaptive immune response. However, in ICI–anti-angiogenic combination therapy, [⁸⁹Zr]Zr-DFO–anti-CD8 Mb PET should be interpreted within the broader context of TME remodeling.

PFC-based ¹⁹F MRI visualizes phagocyte-associated tumor responses through phagocytic uptake, tumor trafficking, and local retention of PFC-labeled cells (*40–42*). In contrast to the circulating [⁸⁹Zr]Zr-DFO–anti-CD8 Mb, the ¹⁹F MRI signal depends on the recruitment and accumulation of PFC-labeled phagocytes within the tumor. It is therefore less sensitive to transient changes in vascular permeability or perfusion that can influence tracer uptake. Consistent with this, neither sorafenib-treated group exhibited an early increase in the ¹⁹F signal at TP1. A recent study showed that immunotherapy-activated CD8⁺ T cells promote the CCR5-dependent recruitment and M1-like polarization of TAMs, suggesting that TAM remodeling is a delayed feature of an effective immunotherapy response (*43*). Accordingly, an increase in [⁸⁹Zr]Zr-DFO–anti-CD8 Mb uptake preceded the later increase in the ¹⁹F signal at TP2, illustrating how longitudinal multimodal imaging captures complementary immune processes across distinct phases of the treatment response. The composition and phagocytic capacity of PFC-laden myeloid subsets vary across tissues and experimental conditions (*44–47*), limiting direct attribution of the PFC-derived ¹⁹F signal to a single marker-defined population and highlighting the need for complementary *ex vivo* analyses. *Ex vivo* analyses revealed reduced CD206⁺ TAM infiltration together with an increased CD206⁻/CD206⁺ TAM ratio in tumors of Combo treated mice, indicating reduced M2 dominance within the TAM compartment. Previous studies in the 4T1 tumor model have linked CD206⁺ M2-like macrophages to tumor relapse, metastatic progression, and tumor-promoting activity (*48, 49*). In line with this, the increased ¹⁹F signal and reduced CD206⁺ dominance suggest that ¹⁹F MRI may help capture features of a less immunosuppressive TME in 4T1 tumors under Combo therapy.

Beyond the immune responses captured by [⁸⁹Zr]Zr-DFO–anti-CD8 Mb PET and ¹⁹F MRI, metabolic reprogramming is another key component of TME remodeling. In TNBC, [¹⁸F]FDG PET has demonstrated value for prognostic stratification and early prediction of response to neoadjuvant chemotherapy (*50, 51*). However, during ICI treatment, its specificity is limited by the inability to distinguish tumor cell uptake from the uptake of activated immune cells (*52, 53*). This was also reflected in our study, where the superior therapeutic response achieved upon Combo treatment was not accompanied by a significant difference in [¹⁸F]FDG uptake. Likewise, [¹⁸F]FDG PET failed to distinguish ICI responders from nonresponders in another TNBC model, whereas granzyme B-targeted PET was qualified to predict response to therapy (*54*). Whole-tumor [¹⁸F]FDG uptake may also be influenced by intratumoral tissue heterogeneity, including necrotic, hypoxic, and poorly perfused regions, which may complicate the interpretation of treatment-related metabolic changes (*55*). Although maximum [¹⁸F]FDG uptake and MTV have been proposed as complementary quantitative metrics (*56, 57*), neither improved discrimination of treatment response in our study. HP MRS, which captures aspects of downstream glycolytic metabolism, showed Lac/Pyr ratio changes that appeared more closely associated with ICI treatment exposure. This interpretation is supported by previous HP MRS studies reporting ICI-associated Lac/Pyr changes across multiple tumor models, although their relationship with treatment response remained inconsistent (*24, 58*). On the one hand, this pattern is plausible, as ICI treatment restores glycolytic activity in reactivated T cells, potentially adding an immune-cell-derived component to lactate production within the TME (*59*). On the other hand, Sorafenib showed a transient increase in both [¹⁸F]FDG uptake and Lac/Pyr ratio at TP1, followed by a decline at TP2. In contrast, previous [¹⁸F]FDG PET studies of high-dose sorafenib therapy in immunodeficient xenograft models showed an early decrease in [¹⁸F]FDG uptake, suggesting that the metabolic effects of sorafenib are highly context dependent, varying with dose, tumor model, immune status, and imaging time point (*60*). These findings suggest that metabolic imaging may primarily capture treatment-associated metabolic changes, particularly during ICI exposure, rather than directly reflecting therapeutic efficacy.

From a translational perspective, this study has relevance for both therapeutic strategy development and response monitoring. Therapeutically, the ICIs–Sorafenib combination builds on clinically available agents. Although sorafenib monotherapy has shown limited activity in metastatic TNBC, later clinical efforts have focused on combination regimens, including chemotherapy-based strategies in HER2-negative or metastatic TNBC, with variable benefit and toxicity concerns (*61–63*). Our results support continued investigation of sorafenib–ICI combinations, in which vascular modulation may reshape the TME to facilitate immune cell infiltration and antitumor immunity. Our imaging strategy also has translational relevance for response monitoring. [¹⁸F]FDG PET is clinically well-established. As the [^89^Zr]Zr-DFO–anti-CD8 Mb used in this study serves as the murine surrogate of the clinical [^89^Zr]Zr-Df-IAB22M2C Mb (*64, 65*), the longitudinal [^89^Zr]Zr-DFO–anti-CD8 Mb PET findings reported here have clear translational relevance. HP MRS is also being evaluated in clinical studies (*66, 67*). ¹⁹F MRI has also shown initial clinical feasibility for cell tracking (*68*), although its application to tumor-associated phagocyte imaging remained largely preclinical. However, the principal translational implication of our study is not the superiority of any single imaging modality, but the value of integrating complementary biomarkers to capture coordinated vascular, immune, and metabolic remodeling during therapy. By combining longitudinal *in vivo* imaging with *ex vivo* validation, we establish a clinically translatable framework for monitoring ICI-sorafenib-based combination therapies.

We acknowledge some limitations of the present study. First, our design was built to detect coordinated vascular, immune, and metabolic remodeling during therapy rather than to resolve the causal hierarchy among them. Establishing directionality will require spatially resolved approaches, such as multiplexed tissue profiling and spatial transcriptomics, which can be readily incorporated into our framework. Second, imaging modalities were acquired in parallel cohorts, a deliberate choice dictated by scanner-specific acquisition requirements and 3R animal-welfare principles. Future integrated PET/MRI protocols may facilitate individual multimodal imaging under a more comparable TME (*69*). Third, our study focused on the 4T1 model, a widely used preclinical model that recapitulates key features of aggressive human TNBC (*70*). Extending this framework across additional cancer types is the logical next step toward establishing generalizability.

In conclusion, combining ICIs with sorafenib enhanced therapeutic efficacy and induced coordinated vascular, immune, and metabolic remodeling in the 4T1 TNBC model. Rather than being defined by a single biomarker, treatment response emerged from the coordinated evolution of these biological processes, creating a TME more permissive to antitumor immunity. These findings highlight the value of longitudinal multimodal imaging for capturing the complexity of ICI-based combination therapy and provide a framework for clinically translatable response monitoring.

## MATERIALS AND METHODS

### Study design

The aim of this study was to determine whether Combo could delay tumor growth in an orthotopic 4T1 TNBC model and to characterize the associated vascular, immune, phagocyte, and metabolic changes using longitudinal multimodal imaging and *ex vivo* validation. Female BALB/c mice bearing orthotopic 4T1 tumors were used for all therapeutic experiments. Mice with size-matched tumors were randomly assigned to four treatment groups: Control, ICIs, Sorafenib, or Combo. Control mice received isotype-matched IgG mAb; mice in the ICIs group received dual anti-PD-1/CTLA-4 mAbs; mice in the Sorafenib group received daily sorafenib; and mice in the Combo group received dual anti-PD-1/CTLA-4 mAbs plus sorafenib. Tumor size was monitored by caliper measurements for treatment allocation and longitudinal growth assessment. Longitudinal imaging was performed using separate cohorts for each imaging modality to reduce cumulative procedural burden associated with repeated anesthesia and tracer administration. [¹⁸F]FDG PET, HP MRS, and ¹⁹F MRI were acquired according to the TP0/TP1/TP2 schedule shown in Fig. 1A, whereas [⁸⁹Zr]Zr-DFO–anti-CD8 Mb PET followed the separate single-tracer-injection protocol shown in Fig. 3A. Flow cytometry and IHC were performed at the indicated time points for *ex vivo* validation, using the corresponding terminal cohorts.

### [⁸⁹Zr]Zr-DFO–anti-CD8 Mb PET

For [89Zr]Zr-DFO–anti-CD8 Mb PET, mice received 2 MBq/10 μg [89Zr]Zr-DFO–anti-CD8 Mb intravenously one day before treatment initiation. This tracer, kindly provided by ImaginAb, serves as the murine surrogate of the clinical [89Zr]Zr-Df-IAB22M2C Mb (*64, 65*). Following the single tracer injection, 10-min static PET scans were acquired at 24, 48, 96, and 192h post-injection, corresponding to D0, D1, D3, and D7 after treatment initiation. PET imaging was performed using a dedicated small-animal Inveon microPET scanner, followed by 3D T2-weighted MRI for anatomical reference and PET/MRI co-registration.

### ¹⁹F MRI

For ¹⁹F MRI, mice received the perfluoro-15-crown-5 ether (PFCE) emulsion intravenously 48h before imaging to label phagocytic cells. Imaging was performed using a dual-tuned ¹H/¹⁹F volume coil with an external ¹⁹F reference tube for signal stability and quality control. Axial ¹⁹F images of the tumor region were acquired using a RARE sequence, followed by 3D T2-weighted MRI for anatomical reference.

### HP MRS

For HP MRS, mice were fasted overnight before imaging. Hyperpolarized [1-¹³C]pyruvate was prepared by dynamic nuclear polarization and injected intravenously through a tail-vein catheter. Dynamic non-localized ¹³C spectra were acquired from the tumor region using a ¹H/¹³C surface coil, with acquisition initiated immediately before pyruvate injection. For longitudinal comparison, Lac/Pyr ratios were normalized to the mean TP0 value of each treatment group.

### [^18^F]FDG PET

For [¹⁸F]FDG PET, mice were fasted overnight before imaging. FDG was injected intravenously, and static PET scans were acquired 60min after injection, followed by 3D T2-weighted MRI for anatomical reference and PET/MRI co-registration.

### Image analysis

PET images were reconstructed using OSEM-3D with Gaussian filtering and registered to anatomical 3D T2-weighted MRI using Inveon Research Workplace. Tumors were manually delineated on MR images, and PET tracer uptake was calculated as %ID/mL after decay correction and normalization to injected activity.

For [⁸⁹Zr]Zr-DFO–anti-CD8 Mb PET radial profile analysis, MR-defined tumor voxels were assigned a normalized distance from the tumor boundary to the tumor center. Mean PET tracer uptake was calculated across radial bins, and center/periphery ratios were derived by splitting voxels at the median normalized radial distance.

¹⁹F MRI data were co-registered with ¹H images and quantified according to previously described methods (*71, 72*). HP MRS data were processed in jMRUI using a model-free Lac/Pyr ratio calculated from the AUC of dynamic ¹³C-lactate and ¹³C-pyruvate signals.

### Flow cytometry

Tumor tissues were processed into single-cell suspensions and stained for flow cytometry. Approximately 2 × 10⁶ cells were stained per sample using the antibody panel listed in table S1. Antibodies were used at titrated concentrations. Samples were acquired on a Cytek Aurora cytometer. When required, flow cytometry files were concatenated using flowCore in R (version 4.5.0). Quality control was performed using flowAI in FCS Express 7, followed by gating and downstream analysis in FCS Express 7. Flow cytometry gating strategies are provided in fig. S2, A and B.

### IHC

Formalin-fixed, paraffin-embedded tumor sections harvested at TP1 were stained for CD8, whereas sections harvested at TP2 were stained for CD31, α-SMA, Iba1, and CD206 to assess vascular and immune features. Quantitative analyses were restricted to viable tumor regions, excluding necrotic areas. CD31⁺ vessels and CD8⁺ cells were manually counted in at least three high-magnification fields per sample. Perivascular α-SMA coverage was quantified from spatially aligned CD31- and α-SMA-stained adjacent sections as the fraction of CD31⁺ area overlapping with α-SMA⁺ signal. CD8⁺ cell distribution was assessed using center/periphery ratios. Iba1 and CD206 staining were quantified in QuPath v0.6.0 as the percentage of positively stained area (*73*).

### Radar plot visualization

Radar plots were generated from group-level imaging, flow cytometry, and IHC readouts after z-score normalization across treatment groups. When multiple readouts represented the same biological feature, z-score-normalized values were averaged with equal weighting for visualization. Radar plots were used for qualitative visualization only and were not used for statistical testing or as validated composite scores.

### Statistical analysis

Statistical analyses were performed using GraphPad Prism 10.0. Data are presented as mean±SEM. For comparisons among multiple groups, one-way ANOVA with Tukey’s or Dunnett’s multiple-comparisons test was used, whereas Brown-Forsythe and Welch ANOVA followed by Dunnett’s T3 test were used when variances were unequal. Longitudinal datasets were analyzed by ordinary or repeated-measures two-way ANOVA with appropriate multiple-comparisons correction. Mixed-effects models were used when missing values precluded repeated-measures analysis. Statistical significance is indicated as \**p*<0.05, \*\**p*<0.01, \*\*\**p*<0.001, and \*\*\*\**p*<0.0001.

## List of Supplementary Materials

Materials and Methods

Table S1

Fig. S1 to S4

## Acknowledgments

We thank Rolf Daniels and Denise Steiner from Department of Pharmaceutical Technology, University of Tuebingen for assistance with the emulsification of the PFC. We thank ImaginAb for kindly providing the DFO-anti-CD8 Mb used in this study. OpenAI ChatGPT (GPT-5.5), Anthropic Claude (Opus 4.8), and Grammarly were used during preparation of this work for language editing and for writing and debugging analysis code used in image registration and quantification, flow cytometry data processing, spectral processing, quantitative analyses, and figure generation. No AI tool was used to generate text presented as original scientific content, to create or alter primary data, or to interpret results. All AI-generated code was reviewed, tested, and revised by the authors; the underlying analysis steps are described in Materials and Methods, and the code is available under Data and Materials Availability. The authors take full responsibility for the accuracy, integrity, and originality of this work.

## Funding

We acknowledge the support of the Deutsche Forschungsgemeinschaft (DFG, German Research Foundation; Grant Nos. 516238665 and 527345502), the DFG under Germany’s Excellence Strategy - EXC 2180 – 390900677, the Werner Siemens Foundation, the Alexander von Humboldt Foundation through the Sofja Kovalevskaja Award, the DKTK German Cancer Consortium Innovation Program “HYPERBOLIC”, the China Scholarship Council (CSC), the Junior Fortüne Grant (F1359045), the Spanish Ministry of Science and Innovation (MCIN/AEI; PRE2020-095268; MCIN/AEI/10.13039/501100011033) and the European Social Fund (“ESF Investing in your future”), as well as the Instituto de Salud Carlos III (ISCIII) through project PI23/00671, co-funded by the European Union.

## Author contributions

Conceptualization: AFM, AMS, SXL, MAN

Methodology: SXL, MAN, SP, ABT, AFM, AMS, IGM, DS, MK, IW

Investigation: SXL, MAN, LK, ABT, IGM Visualization: SXL, IGM, ABT

Funding acquisition: AFM, SXL, MAN, ABT

Project administration: AFM, LK, MK, DS, AMS, LQF

Supervision: AFM, MK, DS, AMS, LQF

Writing – original draft: SXL, ABT, IGM

Writing – review & editing: AFM, MK, DS, AMS, LQF, LK, MAN

## Competing interests

The authors declare that they have no competing interests.

## Data and materials availability

Processed data underlying all quantitative analyses and figures are provided as Source Data. Raw imaging datasets, whole-slide immunohistochemistry images, and flow cytometry data are available from the corresponding author upon reasonable request. Analysis code is available at https://github.com/TheMartinslab/ICIs-Sorafenib-Multimodal-Imaging.

## Notes

### Competing Interest Statement

The authors have declared no competing interest.

