## Supporting Materials for "Immune–metabolic PET/MRI uncovers microenvironmental reprogramming under combined immunotherapy and anti-angiogenic therapy"

### Supplementary Materials

#### Materials and Methods

##### Animals and tumor implantation

Female BALB/c mice were purchased from Charles River Laboratories (Sulzfeld, Germany) and used at 6–8 weeks of age. Animals were maintained in individually ventilated cages with food and water available ad libitum. All experiments were conducted in accordance with German federal regulations and approved by the local authorities (Regierungspräsidium Tübingen).

Orthotopic 4T1 tumors were established by injecting 1.5–2 × $10^5$ 4T1 cells in 25 μL PBS into the fourth mammary fat pad of BALB/c mice. Tumor volume was calculated from caliper measurements using the formula (length × $width^2$) × 0.5.

##### Treatment administration and imaging schedule

Mice with size-matched tumors were randomly assigned to Control, ICIs, Sorafenib, or Combo groups. Control mice received intraperitoneal injections of isotype-matched IgG monoclonal antibodies (mAb) at 3–4 days intervals. ICIs-treated mice received intraperitoneal anti-PD-1 and anti-CTLA-4 mAbs at 3–4 days intervals. Sorafenib-treated mice received daily intraperitoneal sorafenib. Combo-treated mice received both ICIs and sorafenib according to the same schedules.

For antibody treatment, the first dose was 500µg, followed by 200µg for subsequent doses. Anti-PD-1 mAb (clone RPMI-14), anti-CTLA-4 mAb (clone 9H10), and isotype-matched IgG mAbs were obtained from BioXCell (West Lebanon, NH, USA). Sorafenib was administered at 10mg/kg (Nexavar; Bayer Schering Healthcare, Leverkusen, Germany).

[$^{18}$F]FDG PET, HP MRS, and $^{19}$F MRI were performed at baseline before treatment initiation (TP0), day 8 after treatment initiation (TP1), and day 12–15 (TP2), as shown in Figure 1A. CD8 PET followed a separate single-injection schedule. Mice received [$^{89}$Zr]Zr-DFO–anti-CD8 Mb one day before treatment initiation, followed by CD8 PET on treatment days D0, D1, D3, and D7, as shown in Figure 3A.

**CD8 PET tracer preparation and image acquisition**

The DFO-conjugated anti-CD8 minibody (Mb) was radiolabeled with 200MBq [$^{89}$Zr]Zr-oxalate (PerkinElmer, Waltham, MA, USA) per mg protein to generate [$^{89}$Zr]Zr-DFO–anti-CD8 Mb. Briefly, the desired activity was neutralized with a 0.45-fold volume of 2M sodium carbonate and buffered with a 5-fold volume of 0.5M ammonium acetate to achieve pH 6.5–7.0. After adding the protein, labeling was allowed to proceed for 40 min at 25°C, after which the reaction was quenched by adding DTPA solution (0.2%, 40μL per mg protein) and incubated for an additional 10min. Incorporation of the radioisotope was confirmed by iTLC analysis. The elution profile of the labeled protein and radiolabeling of the protein peak were confirmed by high-performance size-exclusion chromatography (HPSEC; BioSep SEC-s2000 column, 300 × 7.8 mm; Phenomenex, Torrance, CA, USA). If radiochemical purity was below 90%, the labeled protein was purified using a Bio-Spin 6 desalting column (Bio-Rad, Hercules, CA, USA) according to the manufacturer's instructions to achieve radiochemical purity above 95%.

For CD8 PET, experimental mice were injected intravenously with 2MBq/10μg [$^{89}$Zr]Zr-DFO–anti-CD8 Mb one day before treatment initiation. Mice were anesthetized with 1–2% isoflurane in breathing air. Following the single tracer injection, 10-min static PET scans were acquired at 24, 48, 96, and 192h post-injection, corresponding to treatment days D0, D1, D3, and D7, respectively.

PET scans were performed using a dedicated small-animal Inveon microPET scanner (Siemens Healthineers, Knoxville, TN, USA) with temperature-controlled heating mats. Subsequently, animals were transferred on the same bed to a 7T preclinical MRI scanner (Bruker BioSpec 70/30, Bruker BioSpin, Ettlingen, Germany). A 3D T2-weighted TurboRARE MRI scan was acquired to provide anatomical reference images for PET/MRI co-registration. MRI parameters were: repetition time/effective echo time, 800/35ms; RARE factor, 16; field of view, $64\times32mm^2$; isotropic spatial resolution, $0.25\times0.25\times0.25mm^3$.

**$^{19}$F MRI probe preparation and image acquisition**

For generation of the $^{19}$F-PFCE emulsion, 10% wt/wt PFCE (Fluorochem, Derbyshire, UK) and 4% wt/wt purified egg lecithin E80S (Lipoid, Ludwigshafen, Germany) were mixed in isotonic buffer containing 10mM HEPES and 2.5% glycerol at pH 7.4 to yield a coarse raw emulsion. The raw emulsion was homogenized using a high-pressure EmulsiFlex-C5 homogenizer (75MPa, 10 cycles; Avestin, Mannheim, Germany). A final particle size of 110±5nm was achieved, as determined by dynamic light scattering using a Zetasizer Nano ZS (Malvern Instruments Ltd., Malvern, UK).

The $^{19}$F-PFCE emulsion was administered intravenously 48h before $^{19}$F MRI using an infusion pump at 100µL/min for 5min. Animals were placed in a dual-tuned $^{1}$H/$^{19}$F linear transmit/receive volume coil (inner diameter, 40mm; Bruker BioSpin GmbH, Ettlingen, Germany), together with an external $^{19}$F reference tube for signal stability and quality control. Axial $^{19}$F images of the tumor region were acquired using a RARE sequence. Imaging parameters were: repetition time/echo time, 3000/7.65ms; RARE factor, 17; field of view, $55\times32mm^2$; slice thickness, 2mm; number of averages, 180. A 3D T2-weighted TurboRARE MRI scan was acquired as described above to provide anatomical reference images.

#### HP MRS pyruvate preparation and acquisition

For HP MRS measurements, animals were fasted overnight before imaging. [1-$^{13}$C]pyruvate solutions were hyperpolarized using a dynamic nuclear polarization polarizer (SpinLab, General Electric, Waukesha, WI, USA). C1-labeled pyruvic acid (Cortecnet, Voisins-le-Bretonneux, France) was polarized according to the manufacturer's instructions. After 60-120min, the hyperpolarized sample was rapidly dissolved and neutralized in an aqueous solution containing 40mM Tris, 60mM NaOH, and 0.1g/L $Na_2EDTA$. The hyperpolarized [1-$^{13}$C]pyruvate solution (80mM) was intravenously injected through a tail-vein catheter.

The tumor region was positioned over a $^{1}$H/$^{13}$C surface coil (2cm diameter; Bruker BioSpin GmbH, Ettlingen, Germany). Acquisition was initiated immediately before injection of the hyperpolarized [1-$^{13}$C]pyruvate solution. Dynamic non-localized $^{13}$C spectra were acquired from the tumor region every second for 512s using a single-pulse free-induction decay (FID) sequence. Acquisition parameters were: flip angle, 5°; repetition time, 1000ms; spectral bandwidth, 3019.32Hz; number of repetitions, 512.

#### [$^{18}$F]FDG PET acquisition

For [$^{18}$F]FDG PET, animals were fasted overnight before imaging. FDG was synthesized according to our marketing license. Animals received 11.1-14.8MBq FDG intravenously. Static PET emission scans were acquired for 10min at 60min after FDG injection, followed by a 13min transmission scan for attenuation correction.

**Image analysis**

PET images acquired in list mode were reconstructed using 3D ordered subset expectation maximization (OSEM-3D), followed by Gaussian filtering with a 1.5mm full width at half maximum (FWHM). Images were registered to anatomical 3D T2-weighted TurboRARE MR images using Inveon Research Workplace (Siemens Healthineers, Knoxville, TN, USA). Tumor ROIs and ROIs for organs of interest were manually delineated on the anatomical MR images. Tracer uptake (%ID/mL) was calculated from the Bq/mL signal after decay correction and normalization to injected activity. For visual comparison, signal intensity and color scales were normalized across groups within each imaging session.

For radial profile analysis, each tumor voxel within the MR-defined tumor ROI was assigned a Euclidean distance to the nearest tumor boundary. Distance values were normalized to the maximum intratumoral distance, yielding a normalized radial distance ranging from 0 at the tumor periphery to 1 at the tumor center. Tumor voxels were divided into ten radial bins of equal normalized-distance intervals, and the mean PET tracer uptake was calculated for each bin. For center/periphery analysis, voxels were split by the median normalized radial distance, assigning approximately half of the tumor voxels to the peripheral region and half to the central region. The center/periphery uptake ratio was calculated as the mean tracer uptake in the central region divided by that in the peripheral region.

Maximum [$^{18}$F]FDG uptake was defined as the highest intratumoral voxel value (%ID/mL) within the MR-defined tumor ROI. Metabolic tumor volume (MTV) was derived from the registered PET images using uptake thresholds of 30% and 40% of this maximum value. MTV was calculated by summing the volumes of voxels exceeding each threshold.

$^{19}F/^{1}H$ MRI images were co-registered and analyzed using Inveon Research Workplace. Quantification of the $^{19}F$ signal followed previously described methods *(1, 2)*.

HP MRS data were processed in jMRUI v5. Manual preprocessing included zero-order phase correction and application of an apodization filter. Subsequent analysis was performed using a model-free approach based on the Lac/Pyr ratio, calculated from the area under the curve (AUC) of the dynamic $^{13}C$-lactate and $^{13}C$-pyruvate signals.

**Flow cytometry sample preparation**

Tumors were excised and digested at 37°C for 30min on a shaker in RPMI containing 2% FCS, 1mg/mL collagenase (Sigma-Aldrich, St. Louis, MO, USA), and 0.1mg/mL DNase I (Sigma-Aldrich, St. Louis, MO, USA). Single-cell suspensions were obtained by sequential filtration through 70µm and 40µm cell strainers. Red blood cells were lysed using ACK lysis buffer (Gibco Life Technologies, Carlsbad, CA, USA) at room temperature. Antibody staining was performed in the presence of anti-mouse CD16/CD32 Fc block (BD Biosciences, San Jose, CA, USA).

**Immunohistochemistry (IHC) staining and quantification**

IHC was performed using an automated immunostainer (Ventana Medical Systems, Tucson, AZ, USA) according to the manufacturer's protocols for open procedures, with minor modifications. Primary antibodies against CD31 (Abcam, Cambridge, UK), α-SMA (Abcam, Cambridge, UK), CD8 (Thermo Fisher Scientific, Waltham, MA, USA), Iba1 (Abcam, Cambridge, UK), and CD206 (Abcam, Cambridge, UK) were used, with appropriate positive and negative controls included. Slides were scanned using a Ventana DP200 system (Roche, Basel, Switzerland), and staining was evaluated by a board-certified pathologist (L.Q.-M.) using Image Viewer software. Quantitative analyses were restricted to viable tumor regions, excluding necrotic areas.

CD31$^{+}$ vessels were counted in at least three randomly selected high-magnification fields per sample to determine microvessel density. For vascular maturity assessment, pathologist-selected regions from the CD31- and α-SMA-stained adjacent sections were spatially aligned using rigid registration in QuPath v0.6.0 *(3, 4)*. Spatially overlapping CD31$^{+}$ and α-SMA$^{+}$ regions were identified, and perivascular α-SMA coverage was quantified as the fraction of CD31$^{+}$ area overlapping with α-SMA$^{+}$ signal.

CD8$^{+}$ cells were manually counted in at least three high-magnification fields per sample and reported as cells per field. For spatial assessment of CD8$^{+}$ cell distribution, tumor regions were classified as center or periphery, and the center/periphery CD8$^{+}$ cell ratio was calculated. Iba1 and CD206 staining was quantified in QuPath v0.6.0 as the percentage of positively stained area in at least three high-magnification fields per sample.

**Radar plot visualization**

Radar plot axes included vessel maturity, CD8 PET uptake, CD8 central enrichment, $^{19}$F phagocyte signal, TAM phenotype shift, glucose uptake, and lactate production.

Vessel maturity was summarized from CD31$^{+}$ vessel density and perivascular α-SMA coverage, with CD31$^{+}$ vessel density inverted so that higher values reflected a more mature vascular phenotype. CD8 PET uptake was represented by tumor tracer uptake on CD8 PET. CD8 central enrichment was summarized from center/periphery ratios derived from CD8 PET and CD8 IHC. $^{19}$F phagocyte signal was represented by tumor $^{19}$F MRI signal. TAM phenotype shift was summarized from CD206 IHC signal and the CD206$^{-}$/CD206$^{+}$ TAM ratio determined by flow cytometry, with CD206 IHC signal inverted so that higher values reflected a shift away from a $CD206^{+}$ phenotype. Glucose uptake was represented by FDG uptake, and lactate production was represented by the Lac/Pyr ratio derived from HP MRS.

For each result, group-level values were z-score normalized across treatment groups. When multiple readouts represented the same biological feature, z-score-normalized values were averaged with equal weighting for visualization. Radar plots were used for qualitative integrative visualization only and were not used for statistical testing or as validated composite scores.

**Table S1. Antibodies used for flow cytometry**

| Marker | Flurophore | Clone | Manufacturer | Catalog No. | Dilution |
|---|---|---|---|---|---|
| CD45.2 | BB515 | 30-F11 | BD Biosciences | 564590 | 1:400 |
| CTLA-4 | PE-CF594 | UC10-4F10-11 | BD Biosciences | 564332 | 1:200 |
| F4/80 | APC-R700 | T45-2342 | BD Biosciences | 565787 | 1:200 |
| MHCII | BUV805 | M5/114.15.2 | BD Biosciences | 748844 | 1:400 |
| CD62L | BUV395 | MEL-14 | BD Biosciences | 740218 | 1:100 |
| CD25 | BV480 | PC61 | BD Biosciences | 566120 | 1:100 |
| CD44 | BUV496 | IM7 | BD Biosciences | 741057 | 1:100 |
| CD3 | BUV737 | 145-2C11 | BD Biosciences | 612771 | 1:200 |
| CD8 | SparkViolet538 | QA17A07 | BioLegend | 155020 | 1:200 |
| Ly6C | APC-Fire750 | HK1.4 | BioLegend | 128046 | 1:100 |
| CD206 | BV421 | C068C2 | BioLegend | 141717 | 1:50 |
| LAG-3 | APC | C9B7W | BioLegend | 125210 | 1:50 |
| TIM-3 | PE-Fire810 | RMT3-23 | BioLegend | 119745 | 1:50 |
| NKp46 | PE | 29A1.4 | BioLegend | 137604 | 1:200 |
| PD-1 | PE-Cy5 | 29F.1A12 | BioLegend | 135255 | 1:50 |
| CD69 | BV650 | H1.2F3 | BioLegend | 104541 | 1:100 |
| PD-L1 | PE-Cy7 | 10F.9G2 | Biolegend | 124314 | 1:50 |
| CD127 | BV785 | A7R34 | BioLegend | 135037 | 1:100 |
| CD11c | BV711 | N418 | BioLegend | 117349 | 1:200 |
| CD11b | AF647 | M1/70 | BioLegend | 101218 | 1:400 |
| B220 | APC-Fire810 | RA3-6B2 | BioLegend | 103278 | 1:200 |
| CD4 | PerCP | RM4-5 | BioLegend | 100538 | 1:100 |
| Ly6G | VioletFluor450 | 1A8 | Tonbo Bioscience | 75-1276 | 1:100 |
| Live/Dead | Ghost Dye | | Cell Signaling Technology | 80862S | 1:100 |

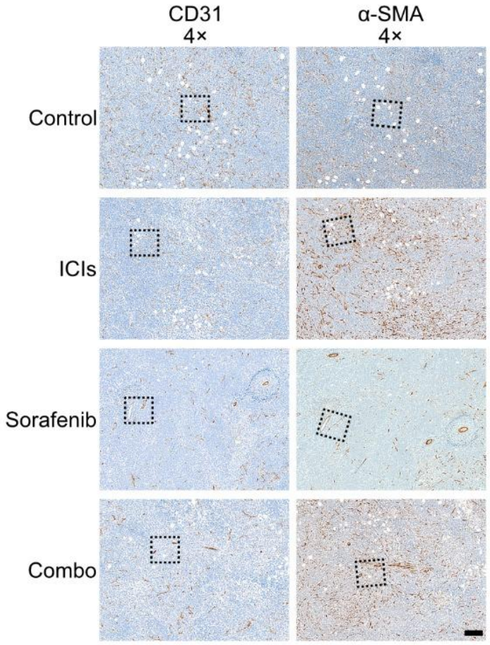


**Supplementary Fig. 1. Low-magnification CD31 and α-SMA staining.**

Representative CD31 and α-SMA IHC images from serial tumor sections of each treatment group shown at low magnification (4×). Scale bars: 1mm.

A

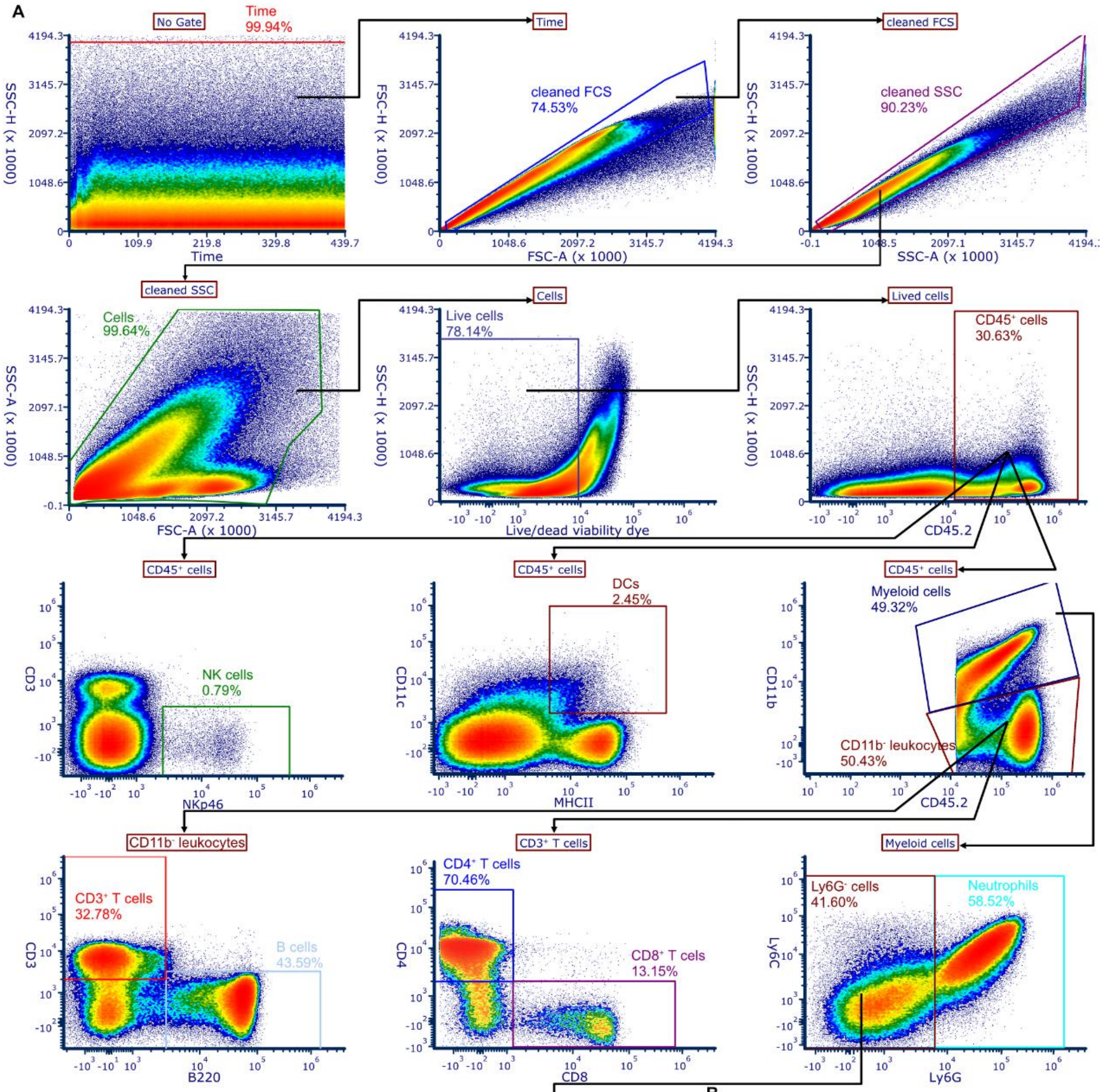


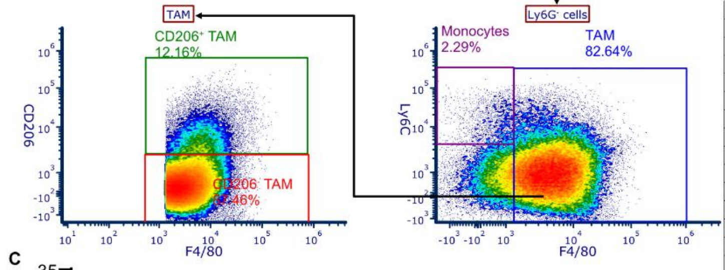


B

| Population | Definition |
| --- | --- |
| CD45$^{+}$ cells | Live CD45$^{+}$ cells |
| Myeloid cells | CD45$^{+}$CD11b$^{+}$ |
| Neutrophils | CD45$^{+}$CD11b$^{+}$Ly6G$^{+}$ |
| Monocytes | CD45$^{+}$CD11b$^{+}$Ly6G$^{-}$Ly6C$^{+}$F4/80$^{-}$ |
| TAMs | CD45$^{+}$CD11b$^{+}$Ly6G$^{-}$F4/80$^{+}$ |
| CD206$^{-}$ TAMs | CD45$^{+}$CD11b$^{+}$Ly6G$^{-}$F4/80$^{+}$CD206$^{-}$ |
| CD206$^{+}$ TAMs | CD45$^{+}$CD11b$^{+}$Ly6G$^{-}$F4/80$^{+}$CD206$^{+}$ |
| Dendritic cells | CD45$^{+}$CD11c$^{+}$MHCII$^{+}$ |
| CD11b$^{-}$ lymphocytes | CD45$^{+}$CD11b$^{-}$ |
| B cells | CD45$^{+}$CD11b$^{-}$CD3$^{-}$B220$^{+}$ |
| CD3$^{+}$ T cells | CD45$^{+}$CD11b$^{-}$CD3$^{+}$ |
| CD4$^{+}$ T cells | CD45$^{+}$CD11b$^{-}$CD3$^{+}$CD4$^{+}$ |
| CD8$^{+}$ T cells | CD45$^{+}$CD11b$^{-}$CD3$^{+}$CD8$^{+}$ |
| NK cells | CD45$^{+}$CD3$^{-}$NKp46$^{+}$ |

C

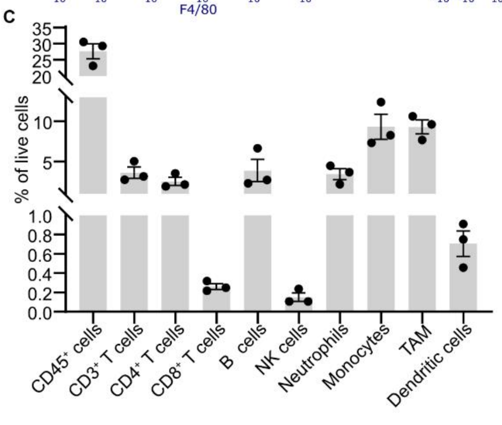

**Supplementary Fig. 2. Flow cytometry gating strategy, immune population definition, and baseline immune profile of untreated 4T1 tumors.**

**(A**) Representative gating strategy for tumor immune-cell profiling. **(B)** Population definitions used for flow cytometry quantification. **(C)** Baseline immune profile of untreated 4T1 tumors, including $CD45^+$ cells, $CD3^+$ T cells, $CD4^+$ T cells, $CD8^+$ T cells, B cells, NK cells, neutrophils, monocytes, tumor-associated macrophages (TAMs), and dendritic cells. Each dot represents one mouse (n = 3).

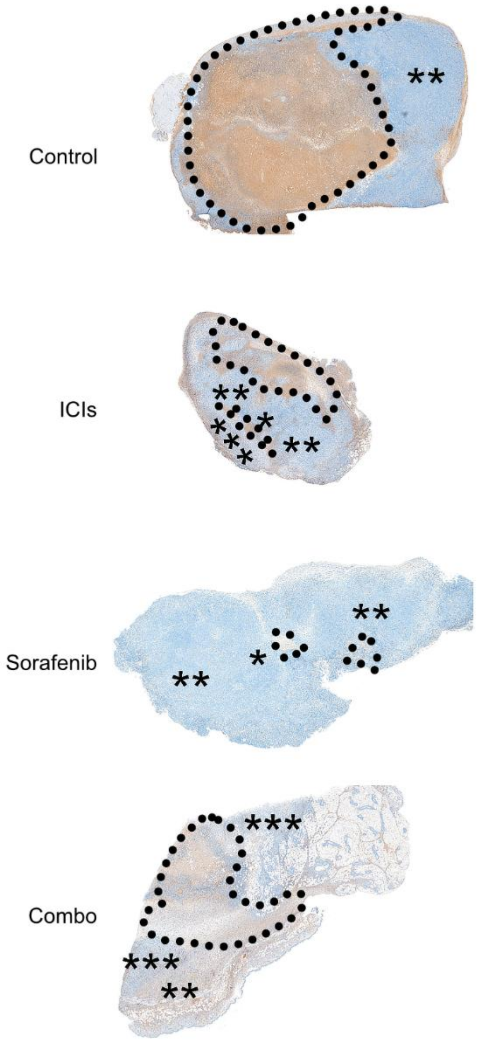


**Supplementary Fig. 3. Low-magnification CD8 staining.**

Representative whole-tumor CD8 IHC images from each treatment group. The dotted outline delineates the necrotic region. *, **, and *** indicate low, intermediate, and high CD8 scores, respectively.

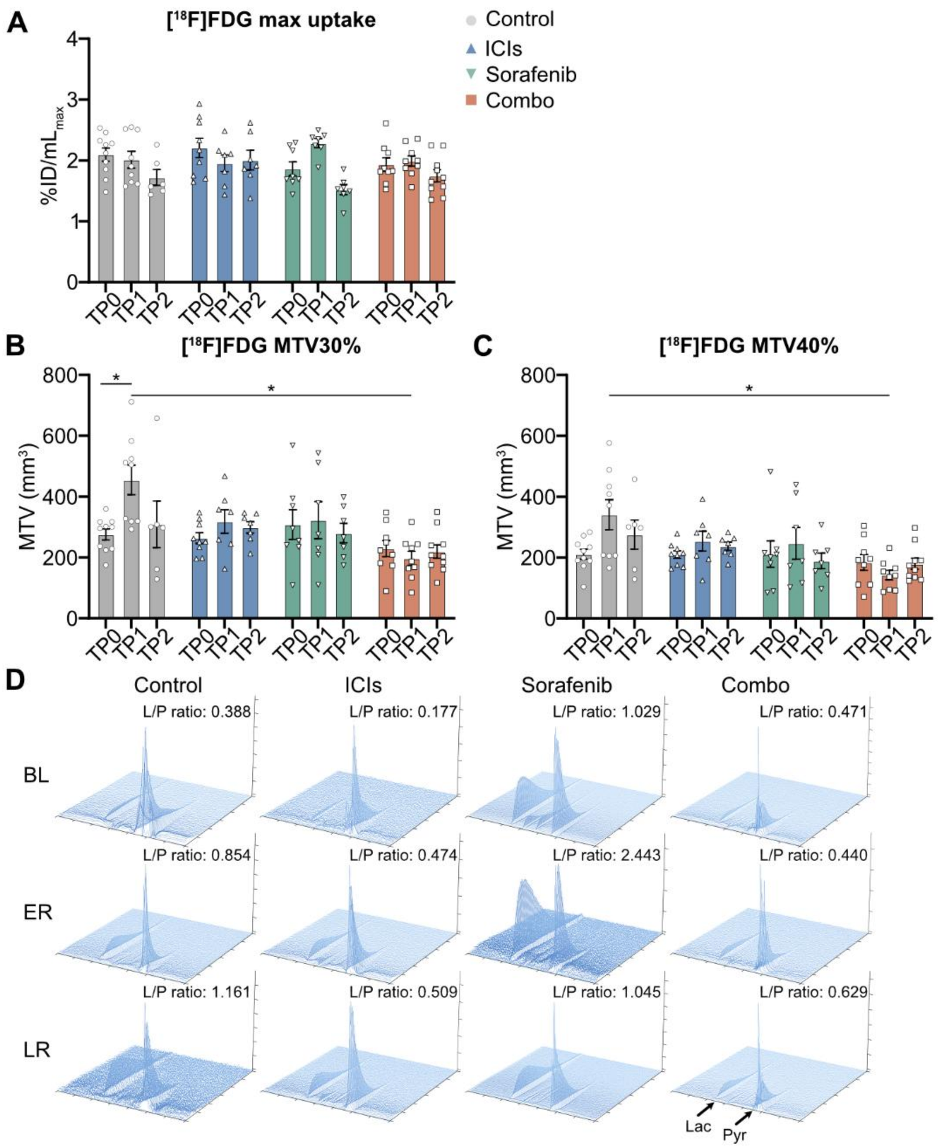


**Supplementary Fig. 4. Additional metabolic imaging analyses.**

**(A)** Longitudinal quantification of maximum [$^{18}$F]FDG uptake as %ID/ml. **(B,C)** Longitudinal quantification of metabolic tumor volume (MTV) using 30% (B) and 40% (C) thresholds, expressed as mm$^3$. **(D)** Representative 3D hyperpolarized $^{13}$C MRS spectra across TP0, TP1, and TP2, providing extended visualization of spectral profiles complementary to Fig.6C. For quantitative panels, n=6–10 mice per group. Each dot represents one mouse. Data are shown as mean±SEM. Statistical analyses were performed using mixed-effects analysis (REML) followed by Tukey's multiple-comparisons test. *$p$<0.05.
